# Novel Mathematical Model Based on Cellular Automata for Study of Alzheimer’s Disease Progress

**DOI:** 10.1101/2022.02.21.481261

**Authors:** Niloofar Jafari, Yashar Sarbaz, Abbas Ebrahimi-kalan, Faegheh Golabi

## Abstract

In recent years, extensive research has been done for the prediction, treatment, and recognition of Alzheimer’s disease (AD). Among these scientific works, mathematical modeling of AD is an efficient way to study the influence of various parameters such as drugs on AD progression. This paper proposes a novel model based on Cellular Automata (CA), a powerful collection of colored cells, for the investigation of AD progress. In our model, the synapses of each neuron have been considered as square cells located around the central cell. The key parameter for the progression of AD in our model is the amount of amyloid-β (Aβ), which is calculated by differential rate equations of the Puri-Li model. Based on the proposed model in this article, we introduce a new definition of AD Rate for a *M* × *L*-neuron network, which can be expanded for the whole space of the hippocampus. To better illustrate the mechanism of this model, we simulate a 3×3 neuron network and discuss the obtained results. Our numerical results show that the variations of some parameters have a great effect on AD progress. For instance, it is obtained that AD Rate is more sensitive to astroglia variations, in comparison to microglia variations. The presented model can improve the scientist's insight into the progress of AD, which will assist them to effectively consider the influence of various parameters on AD.

## 1. Introduction

Nowadays, AD is one of the public neurodegenerative disorders that is primarily caused by the agglomeration of Aβ in the brain (Hardy,Selkoe 2002). It is estimated that AD will affect over 15 million people by 2050. AD is a complicated neurodegenerative disease, which has complex intercellular cross-talks. Many kinds of research have been conducted on cognition, treatment, and reduction of AD progress.

To better understand how AD progresses inside the brain, one of the powerful ways is mathematical modeling. It can help medical society to see the progression of AD, prevent effectively and study the influence of external factors such as drugs on the reduction of AD progress. In the literature, some research articles have been reported for mathematical modeling of AD (Puri,Li 2010; Hao,Friedman 2016; Thuraisingham 2017, 2018; Petrella et al. 2019; Kyrtsos,Baras 2015; Hadjichrysanthou et al. 2018; Dayeh et al. 2018; Jin,Wang 2018; De Caluwé,Dupont 2013; Hoore et al. 2020; Lloret‐Villas et al. 2017; Andrade-Restrepo et al. 2021; Achdou et al. 2013; Helal et al. 2014). The Puri-Li dynamic model (Puri,Li 2010) and its developed mathematical relations (Hao,Friedman 2016) are well-known, familiar papers in the mathematical modeling of AD among scientists. Puri-Li model suggests a dynamic network to show the cross-talk among amyloid-β (Aβ), microglia, neuron, and astroglia, which finally achieves differential equations to be solved numerically. This model has been inspired many researches. For instance, the role of microglia and astroglia in the progression of AD has been investigated based on this model (Thuraisingham 2017). This work has utilized a steady-state population to illustrate that the surviving neurons can be destroyed by microglia and astroglia themselves. In another research, the authors investigated the effect of the Calcium Ion Homeostasis on AD progression by using the Puri-Li model (Thuraisingham 2018).

It is worth mentioning that the mathematical modeling of AD is not limited only to the Puri-Li model. For instance, a system of Smoluchowski equations has been solved by supposing different mechanisms in AD such as neuron-to-neuron prion-like transmission (Bertsch et al. 2017; Franchi et al. 2019). Moreover, a graph theoretical model has been proposed to show the molecular interactions in AD progress (Kyrtsos,Baras 2012). A novel probabilistic computational neurogenetic framework is presented, which contains protein regulatory networks and low-molecular level models (Kasabov et al. 2011).

To the authors’ knowledge, no published work has been reported to analyze and model the AD progress by Cellular Automata (CA). This article aims to propose a neuron-synapse model of AD progression, which will help biologists to schematically observe its progress. The theoretical basis behind this model is the Puri-Li equations for calculating Aβ at a specific time. However, our novelty is the use of CA for representing AD progress. Moreover, the modeling of neuron-to-neuron communications in AD and considering the synapses of a neuron as square cells in CA has been reported for the first time in this paper.

## 2. Analytical Model

In this section, we propose and present an analytical model based on CA for the progress of AD. As explained before, the main reason for the occurrence of AD is the agglomeration of Aβ molecules in the brain. Before embarking on representing of analytical model based on CA, let us review the neuron shape and its structure. Fig. 1 (a) illustrates the various parts of a neuron. Each neuron, which is known as a nerve cell, is composed of three main sections: a cell body, an axon, and dendrites. Here, we only consider the cell body (core), its dendrites, or synapses and ignore other parts such as axon, Myelin sheath, and node of Ranvier (Fig. 1 (b)). This is a realistic assumption because in practice the agglomeration of Aβ happens on synapses. In Fig. 1 (c), the first idea of modeling a neuron and its synapses are shown where the core is located in the center and adjacent square cells are its synapses.

**Fig. 1.**
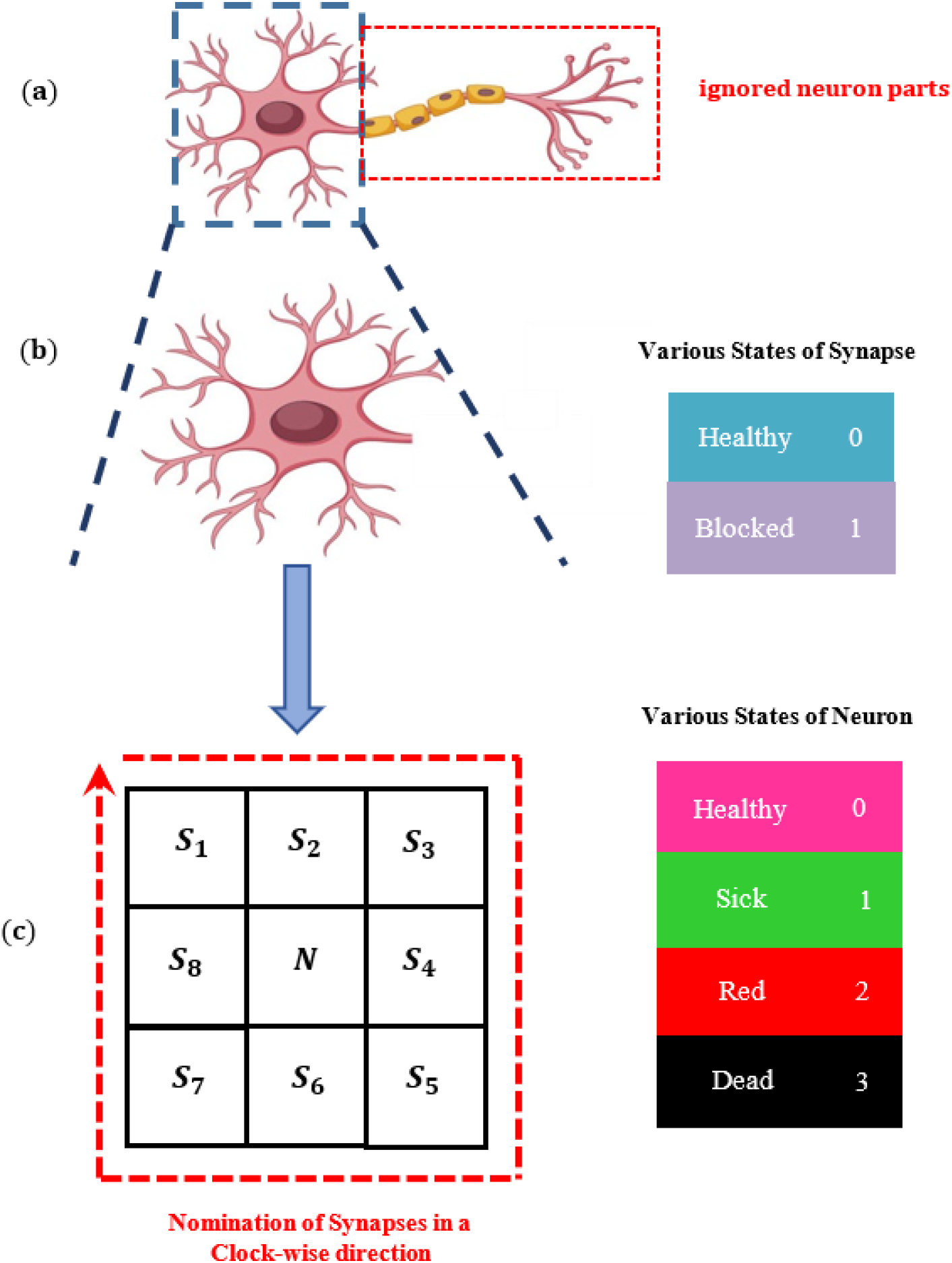
**(a)** Various parts of a neuron, concluding cell body, an axon, and dendrites, **(b)** The cell body (core) and its adjacent synapses, **(c)** CA Network constructed of a central cell for neuron (core) and eight cells for all synapses of it, **(d)** Naming of synapse cells in a clockwise direction.

It should be emphasized that each neuron within the hippocampus has about *synapse*_*total*_ = 1000 synapses. Hence, each synapse cell contains *synapse*_*per cell*_ = *synapse*_*total*_/8 = 125 synapses. Therefore, each square synapse cell (i.e., *S*_1_, *S*_2_, *S*_3_, *S*_4_, *S*_5_, *S*_6_, *S*_7_, *S*_8_ in Fig. 1 (d)) represents 125 synapses. In this paper, we assume that all neurons have equal synapses. The nomination of the synapses is done in a clockwise direction because the aggregation of Aβ molecules is formed continuously (formation of plaques) on synapses. As seen in Fig. 1 (c), each synapse cell (i.e., *S*_1_, *S*_2_, *S*_3_, *S*_4_, *S*_5_, *S*_6_, *S*_7_, *S*_8_ cells) has two states, where the central cell or neuron cell can be in one of 4 states. In what follows, we will explain the various states of synapse cell and neuron more precisely:

### 2.1. Various States of Synapse Cell

As mentioned before, the key parameter for the progression of AD is Aβ molecules, which can be calculated by the Puri-Li dynamic model (Puri,Li 2010):

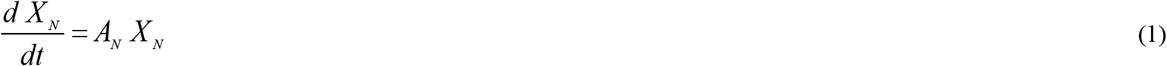

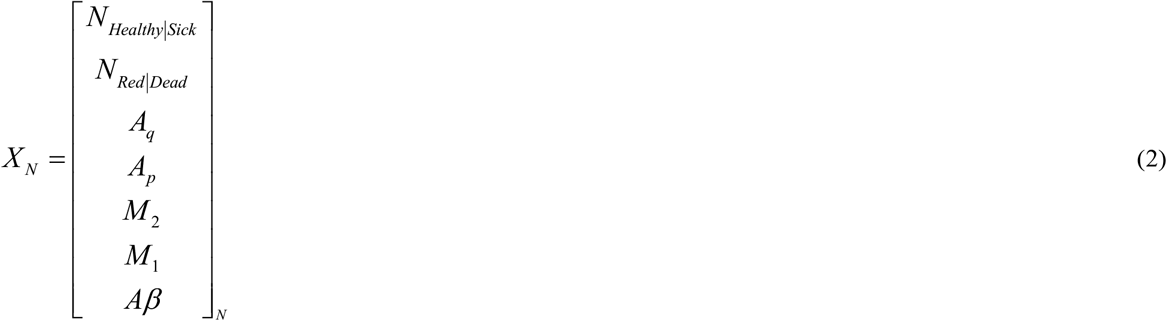

Where

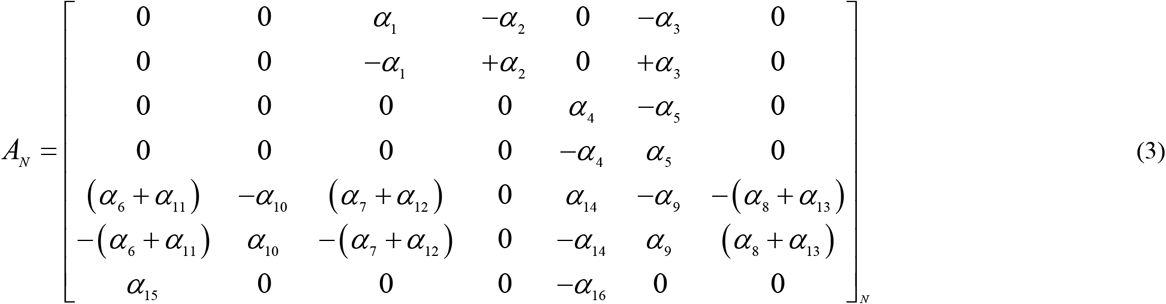

is a parametric matrix, as given in (Puri,Li 2010). In (1)-(3), *N* denotes “Neuron” and *N*_*Healthy*|*Sick*_ and *N*_*Red*|*Dead*_ represent the number of healthy (or sick) and Dead (or red) neurons, respectively. The states of a neuron will be introduced in the rest of the paper. Moreover, in the above relations, *A_q_*, *A_p_*, *M*_2_, *M*_1_, *Aβ* are the number of quiescent astroglia, proliferating astroglia, anti-inflammatory state, pro-inflammatory state, and Aβ molecules, respectively (Puri,Li 2010). The general response of (1) can be expressed as:

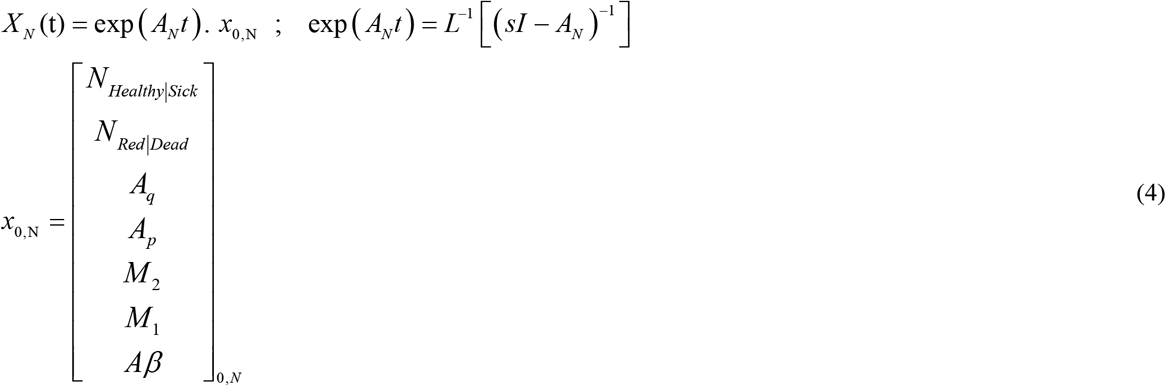

In relation (4), *x*_0,*N*_ is the primary value of each neuron. The parameter of *Aβ* in the above relations determines the whole value of Aβ molecules for all synapses of the neuron. By supposing the time step as *t* = *t*_0_, the blocked synapse cells will be obtained as follows:

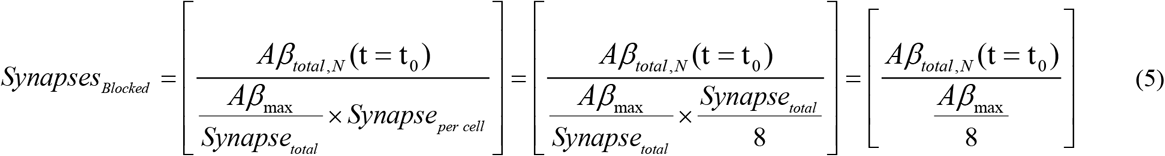

In the above relation, *Aβ*_*total,N*_(*t* = *t*_0_) is the total value of Aβ molecules of the neuron, calculated by the Puri-Li model and *Aβ*_*max*_ is the maximum value of Aβ molecules for a neuron to be blocked completely. Furthermore, the “[]” denotes the *floor* function.

Now, the status of synapse cell can be defined as:

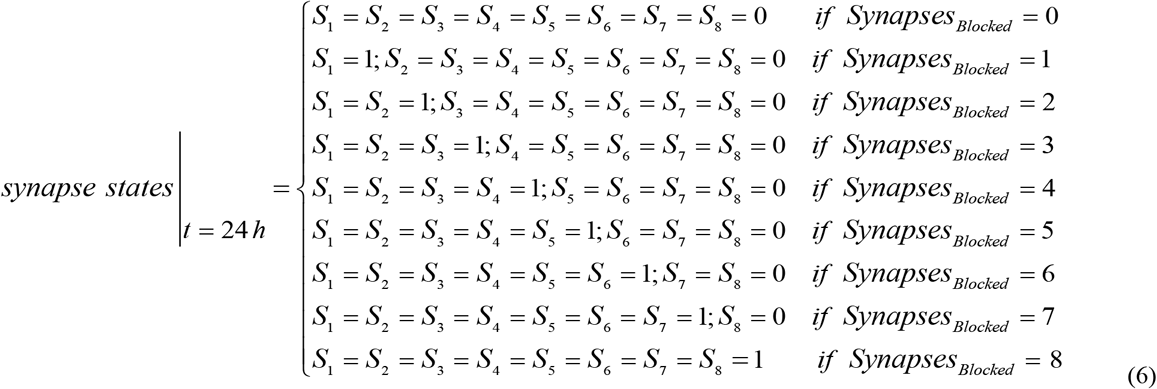

Fig. 2 illustrates the various states of the central neuron and its adjacent synapse cells. To be brief, state “0” in each synapse cell occurs when the amount of Aβ molecules is not sufficient to block the synapses of that cell, while state “1” is related for a case when the whole synapses are blocked by Aβ molecules.

**Fig. 2.**
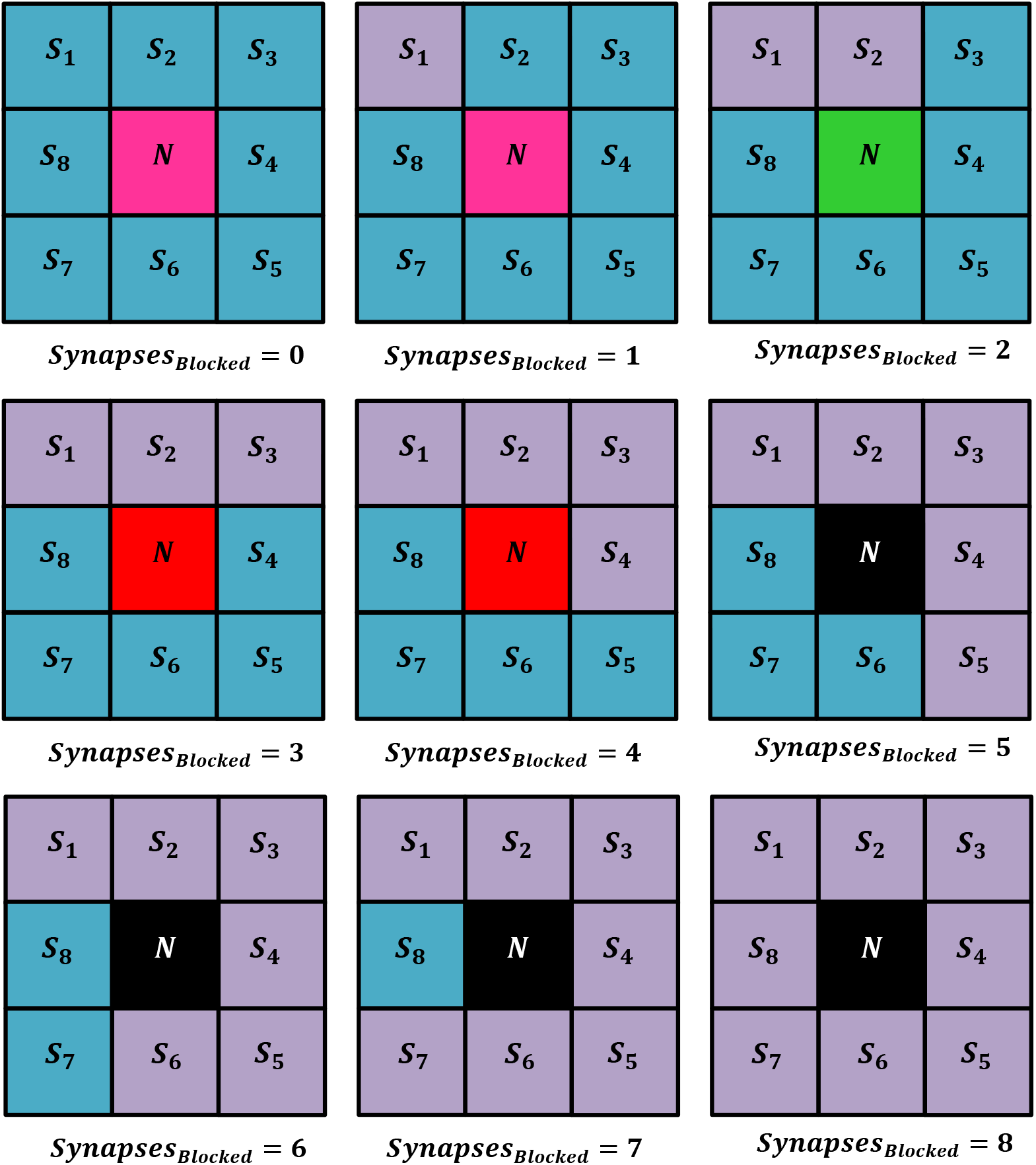
Various states of an isolated neuron and its adjacent synapse cells.

### 2.2. Various States of Neuron

As seen in Fig. 2, each neuron can be in one of the following states, which depends on the states of adjacent synapse cell:

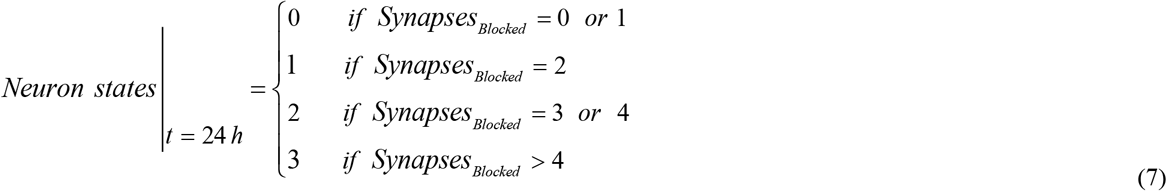

In the literature, the neuron is considered as an “alive (healthy)” or “Dead” neuron. However, since the conversion of the healthy neuron to dead happens gradually, here we define two additional middle states (sick and red states) for a neuron, as follows.

#### 2.2.1. Healthy Neuron (state “0”)

In this state, the amount of Aβ is very low and thus the neuron maintains its connections with other adjacent neurons. Based on our definition, a healthy neuron is a case in which no more than 12.5% of synapse cells are blocked.

#### 2.2.2. Sick Neuron (state “1”)

As defined in (7), a sick neuron is a neuron in which 25% of its synapse are blocked and thus these blocked synapses cannot communicate with other synapses. Biologically, the conversion probability of this state to the Red state is high. In practice, various factors such as age, high cholesterol, diabetes, hypertension, inactivity, and head injury can potentially develop this kind of neuron within the hippocampus.

#### 2.2.3. Red Neuron (state “2”)

In this case, the neuron is in an emergency mode, because it has problems in communicating and sending or receiving signals from other neurons. In our definition, the red state is related to a neuron, which between 25% up to 50% of its synapses are blocked by Aβ molecules. In this state, the neuron is not able to return to a sick state and thus its situation should be stabilized by drugs to prevent its blocking procedure.

#### 2.2.4. Dead Neuron (state “3”)

In this state, more than 50% of synapses have been blocked and hence neuron-to-neuron communication is disrupted. As a neuron dies, it cannot be returned to previous states, and thus some useful methods such as drugs, music, and activity should be performed to prevent this state.

### 2.3. Model of Neuron-to-Neuron Communications in a *M* × *L*–Network

Consider Fig. 3, a *M* × *L*–Network, where each central neuron (denoted by *(i,j)*) is surrounded by 8 synapse cells. In this figure, *i,j* represent the row and column number of *(i,j)*-th neuron, respectively. If we assume that Puri-Li equations can be solved for each neuron in this network with the different primary conditions, then:

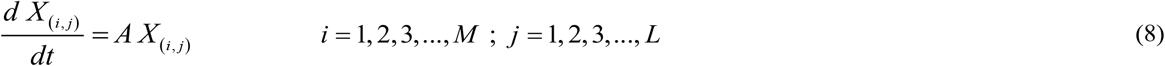

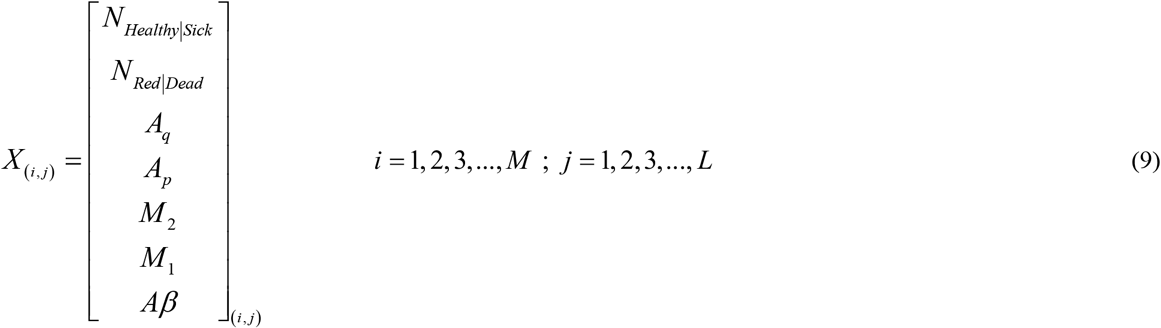

Where

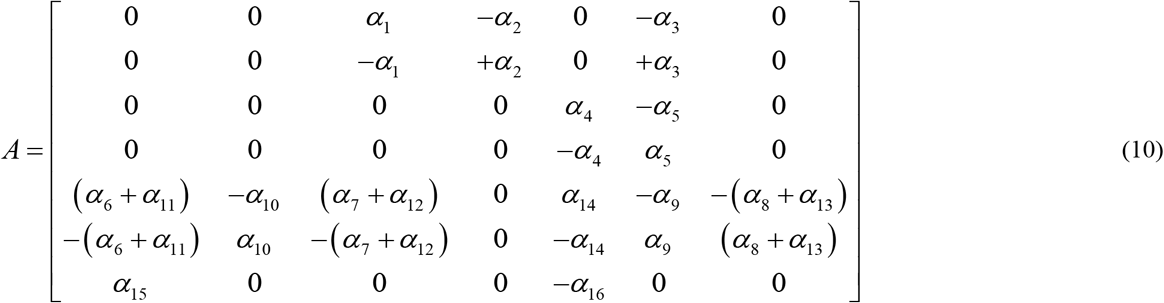

**Fig. 3.**
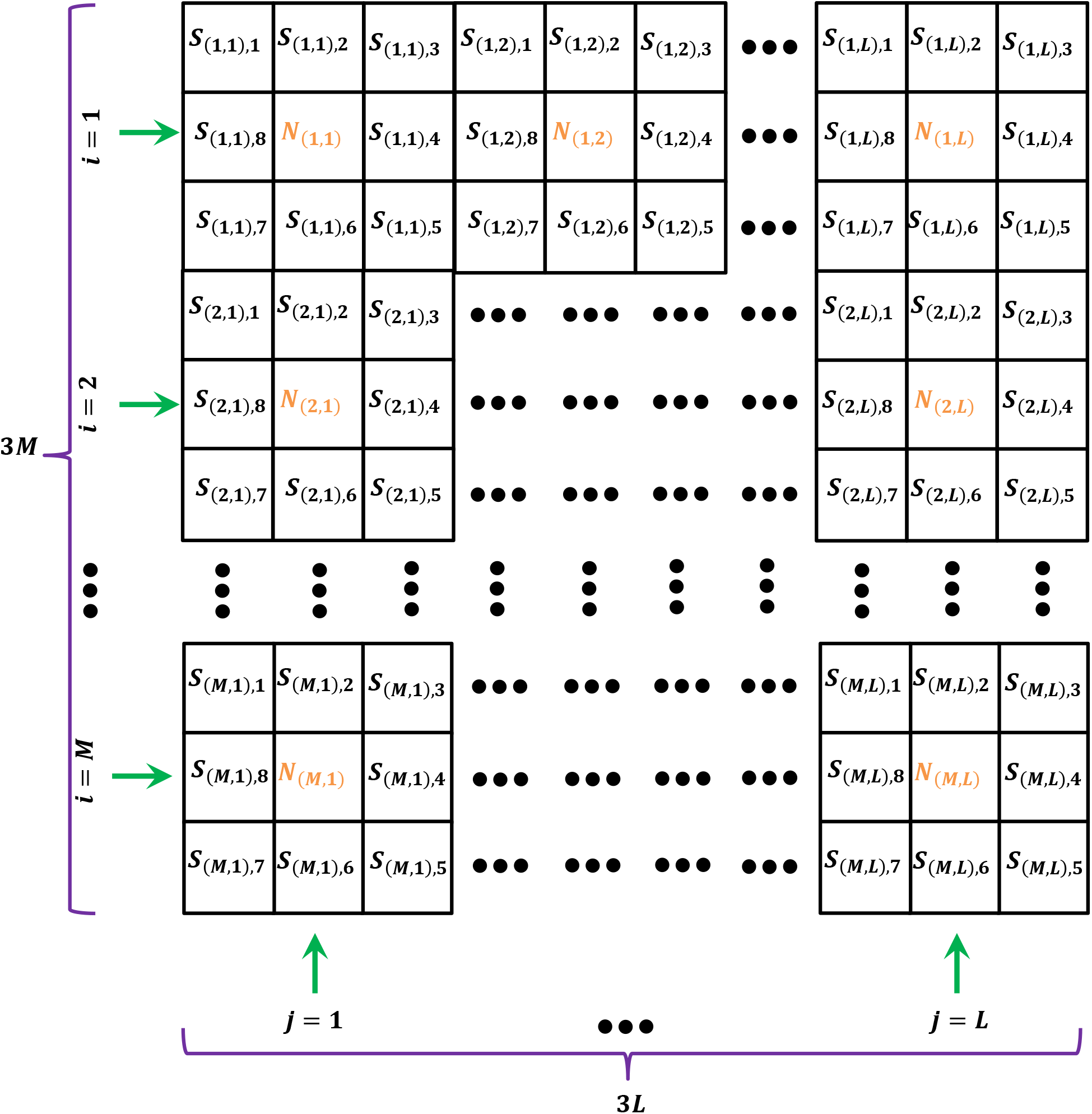
Cellular automata for *M* × *L*– Neuron Network

As explained before, by assuming the time step of *t* = *t*_0_ and beginning with *t* = 0, the general response of (8) for *(i,j)*-th neuron can be written as:

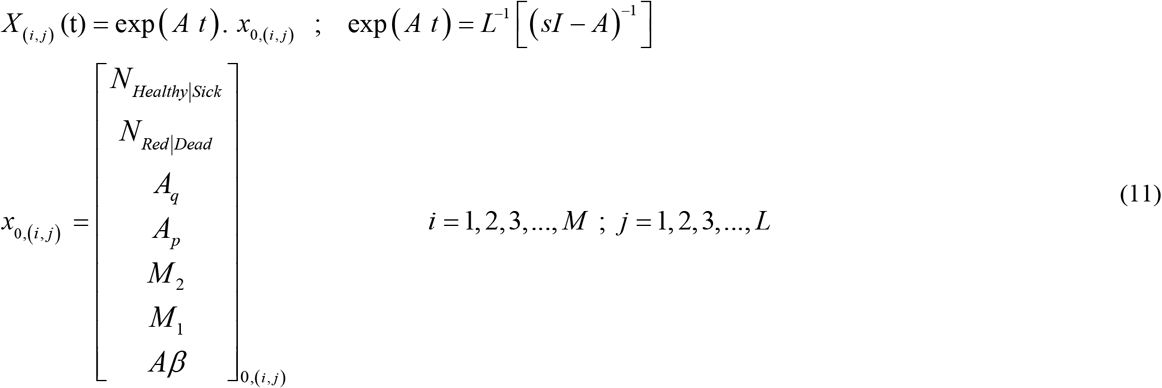

Now, the total numbers of blocked cells at *t* = *t*_0_ can be calculated:

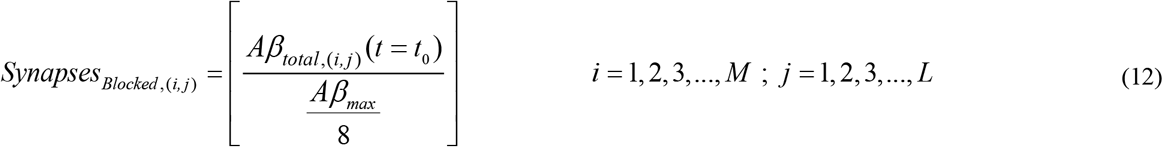

Finally, the synapse cell states for each neuron in *M* × *L*– Neuron Network is derived:

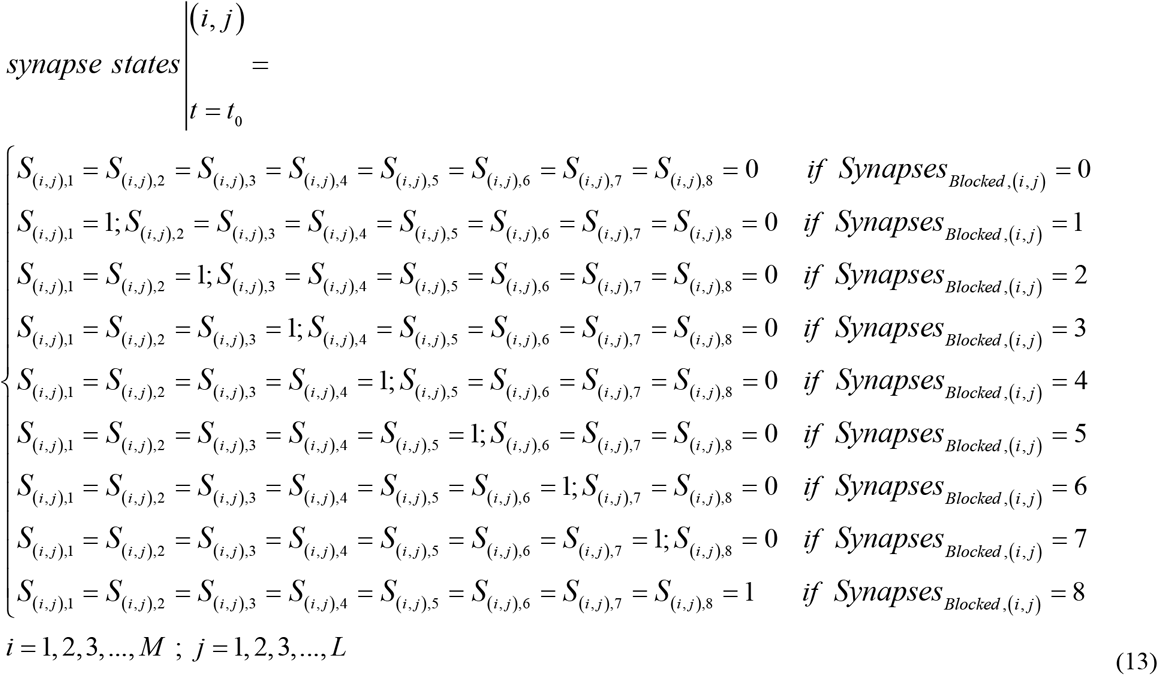

In relation (12), each neuron has been considered as an isolated neuron, as similarly defined in relation (5) of section 2.1 for one neuron (see Fig. 2). To effectively model the neuron-to-neuron communication of synapse cells, the total blocked synapse cells for *(i,j)*-th neuron is determined by the following relation:

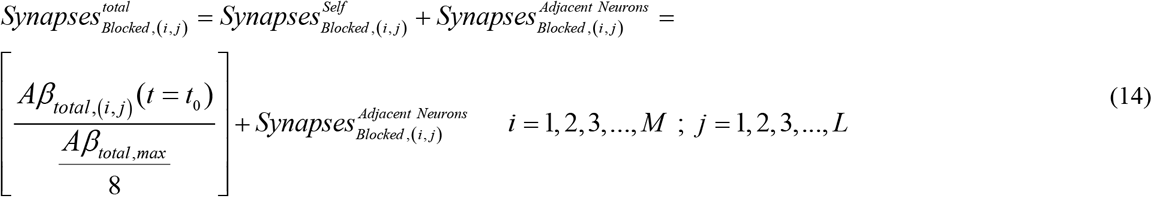

Although the synapses (of a specific neuron) also send and receive signals for further neurons within the brain, however, the modeling of these connections is very complicated and is outside the scope of this article. Therefore, we only consider neuron-to-neuron communication of adjacent synapse cells in *M* × *L*– Neuron Network. By defining (14), let us briefly introduce and review three kinds of neurons in our network: ***1-Corner Neurons*** (i.e., *N*_(1,1)_, *N*_(1,*L*)_, *N*_(*M*,1)_, *N*_(*M*,*L*)_): In a *M* × *L*– Neuron Network, there exist only 4 corner neurons. These neurons have connections with 15 synapse cells. ***2-Row and Column Neurons*** (i.e., *N*_(*i,j*)_, *where i* = 1, *M and j* = 2,3, …, *L* − 1 *or i* = 2,3, …, *M* − 1 *and j* = 1, *L*): the number of these neurons is the *M* × *L*– Neurons Network is 2(*M* + *L*) − 8, which are located in the first and last columns or rows, except corner neurons, and are connected to 19 synapse cells. ***3-Central Neurons:*** In a *M* × *L*– Neurons Network, these neurons have connections with 24 synapse cells and the number of them is *ML* − 2*M* − 2*L* + 4. By defining these categories, neuron states can be updated and modified as follows:

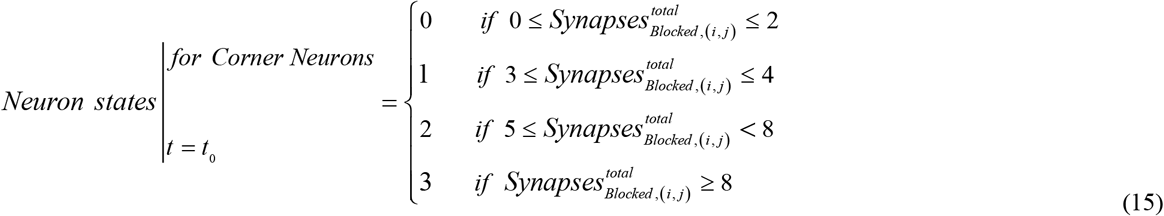

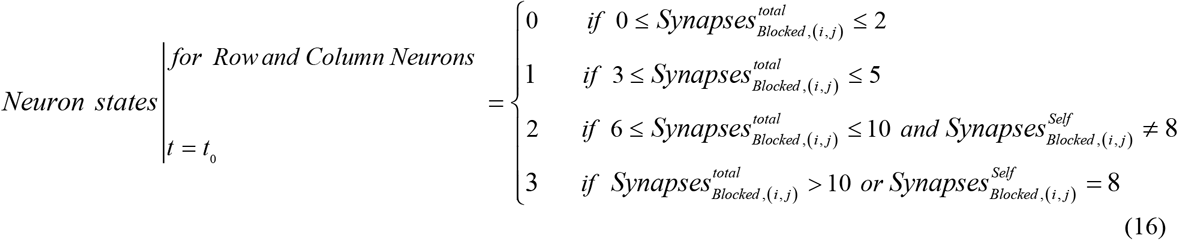

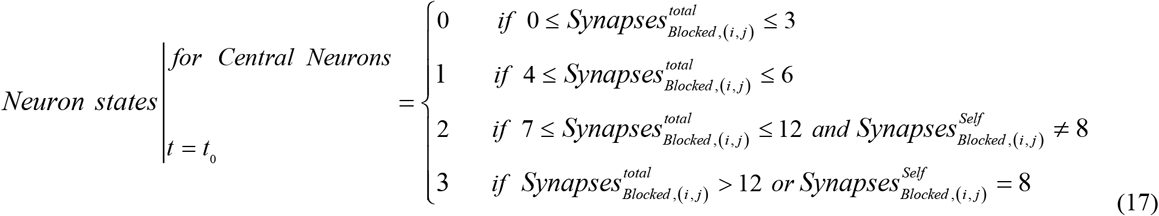

Therefore, by defining neuron states for modeling neuron-to-neuron communications in relations (15)-(17), Fig. 2 should be modified because it has been depicted for an isolated neuron. For instance, all synapse cells for a neuron in a *M* × *L*– Neuron Network can be healthy but its state can be in a sick or red state (because neuron state also depends on the synapse states of adjacent neurons, see relation 14). Accordingly, we can define the Alzheimer’s rate for the progress of this disease. For this purpose, consider a 2 × 2–Network, as shown in Fig. 4. Suppose that 3 neurons have died in this network due to the increment of Aβ molecules (see Fig. 4) and thus the remaining healthy neuron cannot communicate with other neurons. Thus, the alive neuron will die and AD will occur. Therefore, we can define Alzheimer’s Rate as follows:

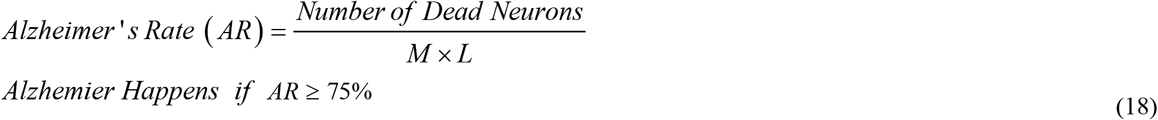

**Fig. 4.**
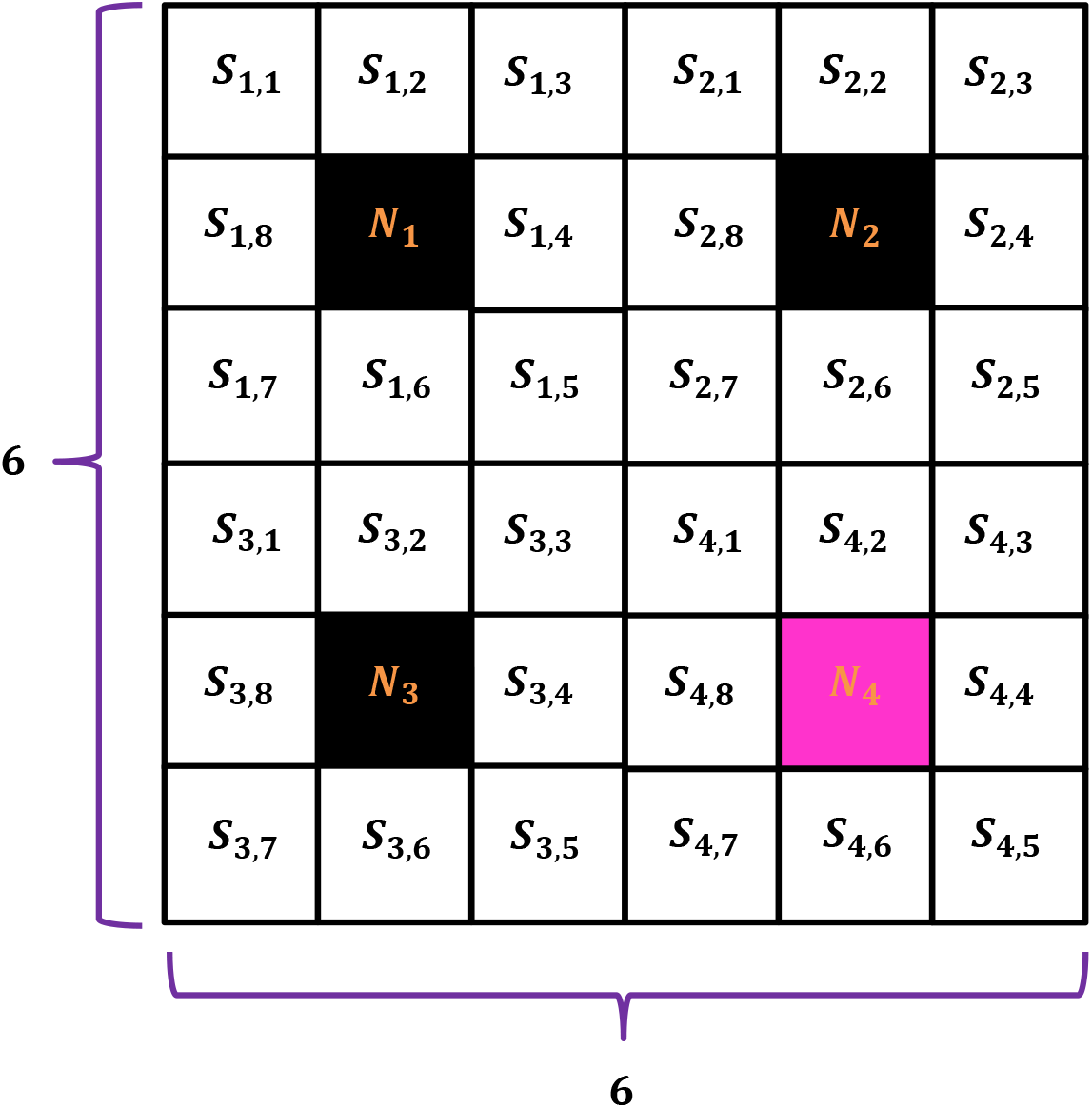
Cellular automata for a 2 × 2–Network, for the definition of Alzheimer’s rate

Similar to the relation (18), Critical and Warning Rates can be defined for a *M* × *L*– Neuron Network:

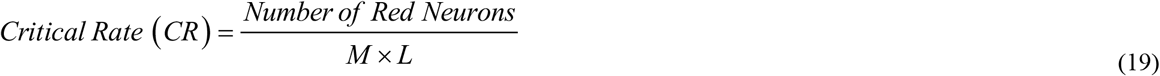

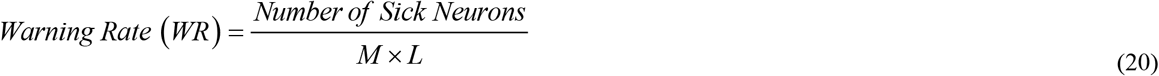

As explained before, the time step is supposed to be *t* = *t*_0_ in Puri-Li equations (see the relations (9)-(11)). By ignoring the internal mechanism within the brain, synapse state for the next time step (*t* = 2*t*_0_) can be computed as follows:

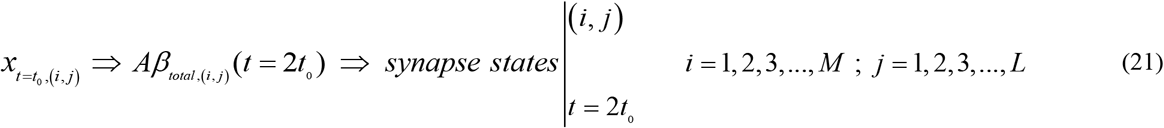

In practice, some internal processes within the hippocampus reduce the amount of produced Aβ molecules, and thus the relation (21) should be modified:

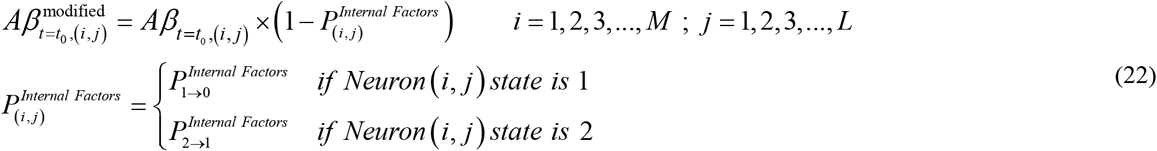

In (22), for instance, 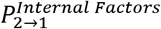 is a probabilistic function for converting the red state (state “2”) of *(i,j)*-th neuron to the sick state (state “1”). It is obvious the as the neuron state goes toward the dead state, the probability decreases:

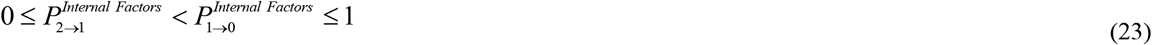

As a result, the primary values for *(i,j)*-th neuron at *t* = *t*_0_ can be expressed as follows:

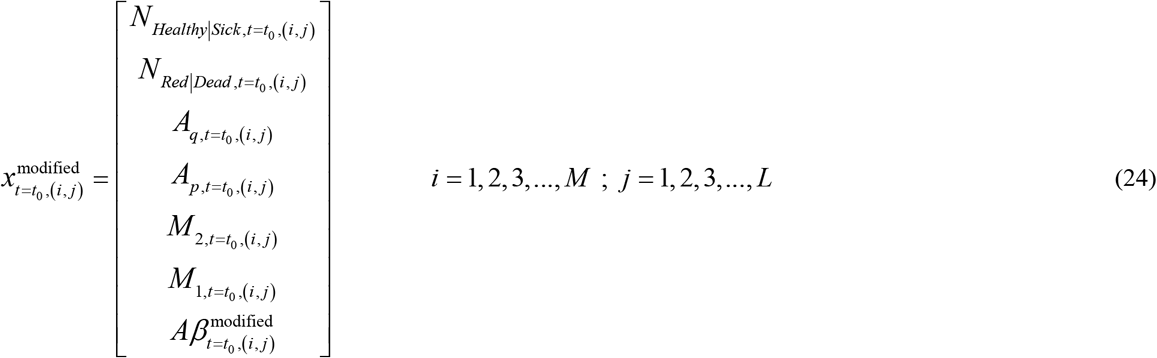

## 3. Results and Discussions

We investigate the numerical results of the proposed model in this section. As we know, AD begins within the hippocampus, which occupies about 2.5 cm^2^ in the two-dimensional surface. By supposing the area of a neuron as 250 μm^2^, then the number of cells will be obtained about 10^6^. The simulation of this number of neuron cells is outside the scope of this research, therefore, we only focus on the CA network with a low number of neurons, to get biological insight into the progression of AD. Consider a 3×3 neuron network with its simulation parameters given in Table. 1. The mathematical parameters of the Puri-Li dynamic model are given in Table. 2. Besides, to solve the state-space equations of (8), the initial values of neurons’ parameters are needed, which are given in Table. 3 for three various random cases.

**Table 1.**
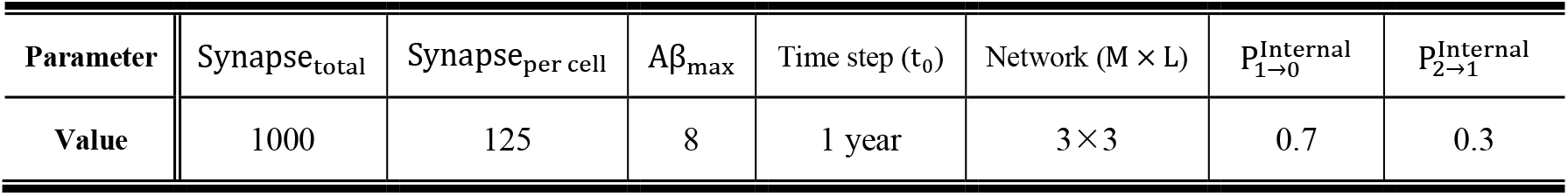
Simulation Parameters of CA Network.

| Parameter | Synapse<sub>total</sub> | Synapse<sub>per cell</sub> | $A\beta_{\max}$ | Time step ($t_0$) | Network ($M \times L$) | $p_{1 \rightarrow 0}^{\text{Internal}}$ | $p_{2 \rightarrow 1}^{\text{Internal}}$ |
| --- | --- | --- | --- | --- | --- | --- | --- |
| Value | 1000 | 125 | 8 | 1 year | $3 \times 3$ | 0.7 | 0.3 |

**Table 2.** Mathematical Parameters of Puri-Li model (1/*year*).

| Parameter | $\alpha_1$ | $\alpha_2$ | $\alpha_3$ | $\alpha_4$ | $\alpha_5$ | $\alpha_6$ | $\alpha_7$ | $\alpha_8$ | $\alpha_9$ | $\alpha_{10}$ | $\alpha_{11}$ | $\alpha_{12}$ | $\alpha_{13}$ | $\alpha_{14}$ | $\alpha_{15}$ | $\alpha_{16}$ |
| --- | --- | --- | --- | --- | --- | --- | --- | --- | --- | --- | --- | --- | --- | --- | --- | --- |
| Value | $10^{-5}$ | $10^{-3}$ | $10^{-2}$ | $10^{-4}$ | $10^{-2}$ | $10^{-2}$ | $10^{-4}$ | $10^{-2}$ | $10^{-2}$ | $10^{-2}$ | $10^{-2}$ | $10^{-4}$ | $10^{-2}$ | $10^{-4}$ | 1 | $10^{-2}$ |

**Table 3.** Initial values of various parameters of neurons in a 3×3 network for three random cases.

| Neuron<br>($i,j$) | Parameters of Case 1 | | | | | Parameters of Case 2 | | | | | Parameters of Case 3 | | | | |
| --- | --- | --- | --- | --- | --- | --- | --- | --- | --- | --- | --- | --- | --- | --- | --- |
| | $A_q$ | $A_p$ | $M_1$ | $M_2$ | $A\beta$ | $A_q$ | $A_p$ | $M_1$ | $M_2$ | $A\beta$ | $A_q$ | $A_p$ | $M_1$ | $M_2$ | $A\beta$ |
| (1,1) | 10 | 0.1 | 0.1 | 11 | 0.2 | 9 | 0.12 | 0.12 | 10 | 0.25 | 10 | 0.12 | 0.1 | 11 | 0.35 |
| (1,2) | 32 | 0.52 | 0.5 | 28 | 3.4 | 10 | 0.13 | 0.1 | 11 | 0.3 | 52 | 0.49 | 0.53 | 39 | 5.2 |
| (1,3) | 10 | 0.1 | 0.11 | 10 | 0.12 | 13 | 0.25 | 0.2 | 12 | 0.8 | 9 | 0.12 | 0.11 | 10 | 0.2 |
| (2,1) | 13 | 0.25 | 0.2 | 12 | 0.8 | 14 | 0.23 | 0.24 | 13.5 | 1.2 | 10 | 0.1 | 0.1 | 10 | 0.18 |
| (2,2) | 14 | 0.23 | 0.22 | 13.2 | 1.1 | 62 | 0.6 | 0.58 | 51 | 6.15 | 10 | 0.11 | 0.12 | 9 | 0.22 |
| (2,3) | 19.7 | 0.45 | 0.41 | 14 | 2.18 | 10 | 0.1 | 0.1 | 10 | 0.18 | 19.1 | 0.43 | 0.35 | 14.9 | 2.6 |
| (3,1) | 43 | 0.57 | 0.51 | 31 | 4.5 | 13 | 0.23 | 0.2 | 13.2 | 1.08 | 14 | 0.23 | 0.24 | 13.5 | 1.2 |
| (3,2) | 19.6 | 0.44 | 0.38 | 13.8 | 2.17 | 19.5 | 0.44 | 0.37 | 13.9 | 2.3 | 43 | 0.57 | 0.51 | 31 | 4.7 |
| (3,3) | 64 | 0.59 | 0.61 | 53 | 6.2 | 9 | 0.13 | 0.1 | 10 | 0.28 | 10 | 0.1 | 0.13 | 9 | 0.21 |

Fig. 5 represents schematically the initial value of neurons and their synapse cells in a 3×3 network at *t* = 0 for each case. As explained before, the state of each neuron in a 3×3 network depends on the states of its synapses and the synapses of adjacent neurons (see the relations (14)-(17)). For instance, consider neuron (2,1) in the first case (see Fig. 5 (a)), which all of its synapses are healthy but its state is sick. The state of this neuron is calculated by the relation (16) and has been affected by some blocked synapses of adjacent neurons (i.e., neuron (3,1) and neuron (3,2)). This phenomenon is consistent with reality because as the synapses of adjacent neurons are blocked by the agglomeration of Aβ molecules, they generate NTF that can convert survived neurons to dead ones.

**Fig. 5.**
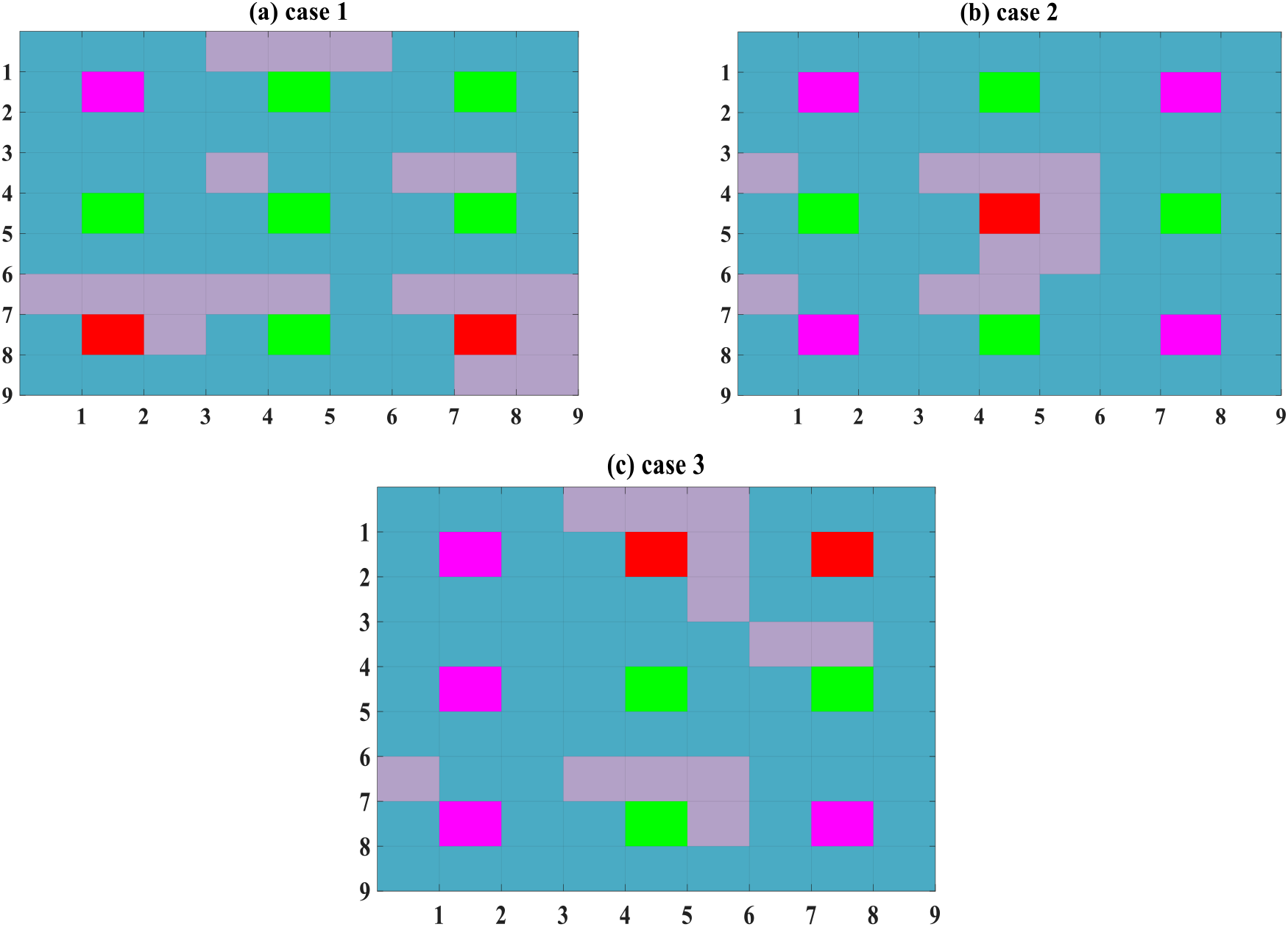
Initial states of neurons and their synapses in a 3×3 network at *t* = 0 for three cases.

Let us see what happens for a 3×3 network after one year. Fig. 6 illustrates the simulation results (provided by MATLAB software) of the CA network at *t* = 1 *year*for three cases. In case 1, all neurons except neuron (1,1) have red states. In this case, AR is zero but CR is about 88.8% which needs special attention. Fortunately, some internal mechanisms within the hippocampus reduce the Aβ agglomeration, and thus the CR decreases. These factors are modeled by probabilistic functions in (22). For the second case, after one year, AR is zero but CR reaches 22.2 % and WR is about 55.5 %, which is a low-risk situation. For the third case, as seen in Fig. 6 (c), CR is similar to the second case (i.e., CR=22.2%) while WR is 66.6%.

**Fig. 6.**
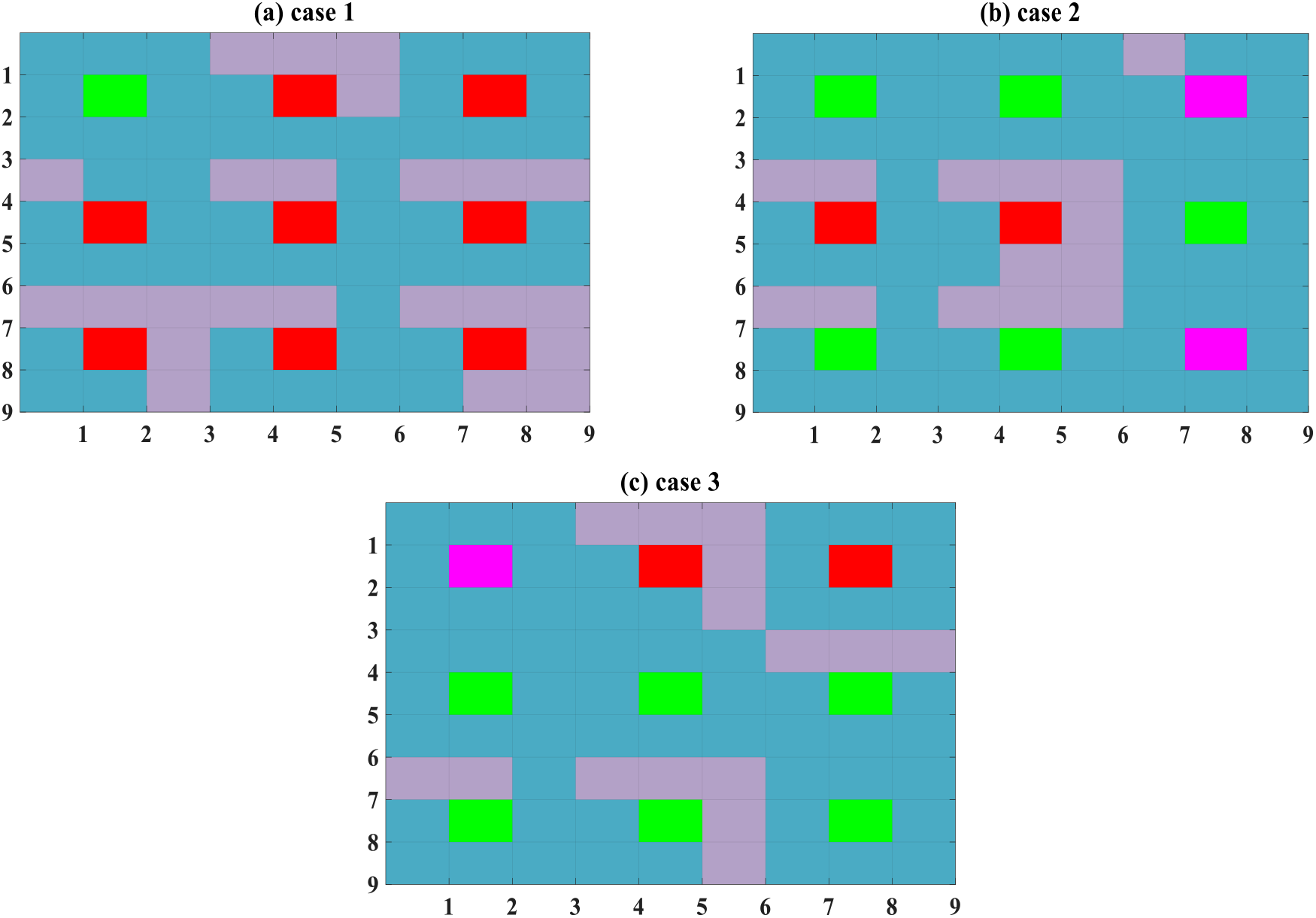
CA model of neurons and their synapses in the 3×3 network at *t* = 1 *year* for three cases.

Fig. 7 represents the progress of AD after two years for three cases. It can be seen from this figure that internal factors have decreased CR for all cases. For the first case, it has reached 11.1 % while it is zero for the second and the third cases. The network has passed from a critical state for the first case while the warning rate (WR) is high (*WR* ≈ 88.8 %). As observed in Fig. 7 (b), (c), the WR for the second and third cases decreases (WR for the second and the third case is 22% and 11%, respectively).

**Fig. 7.**
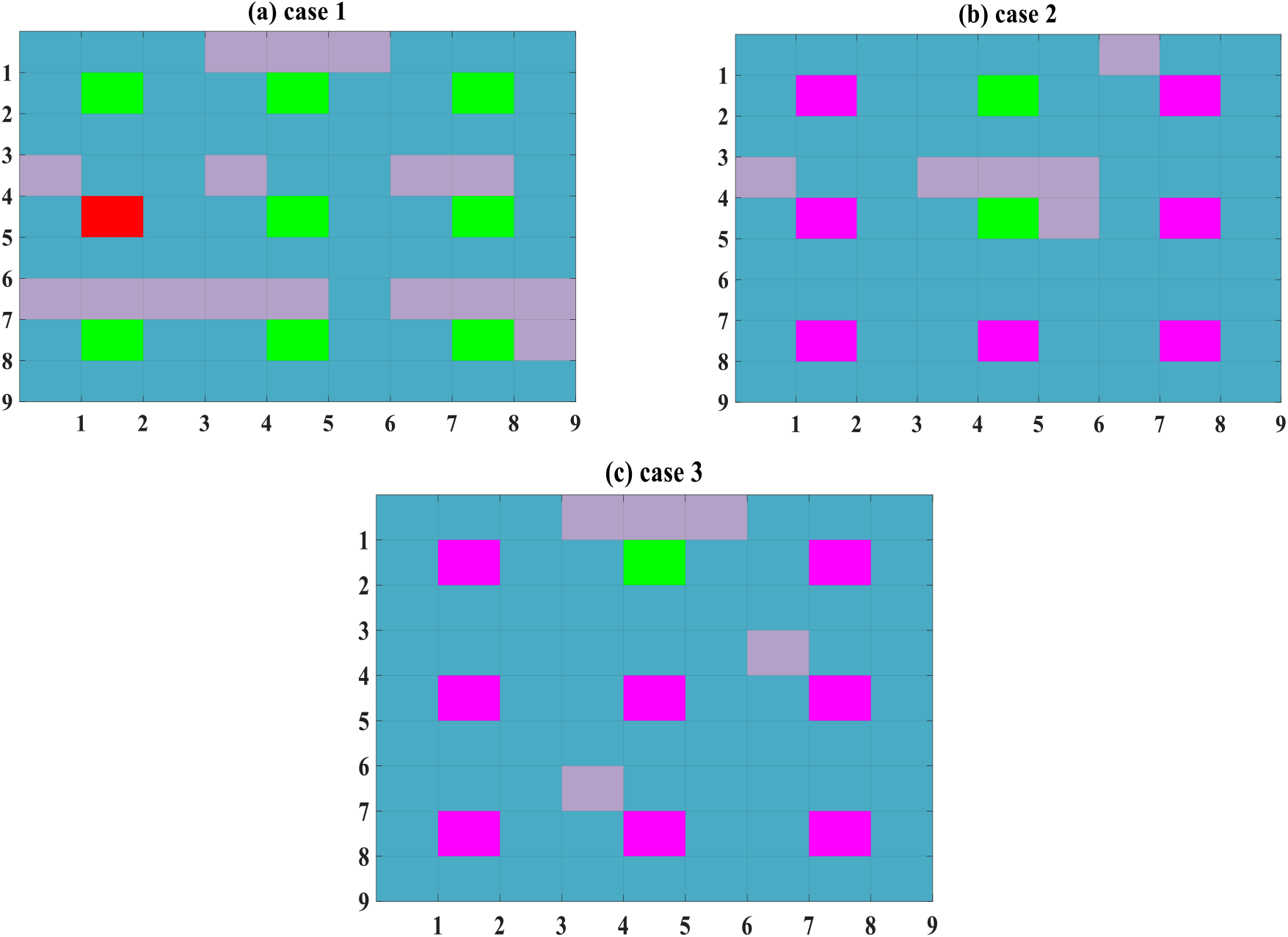
CA model of neurons and their synapses in a 3×3 network at *t* = 2 *year* for three cases.

To investigate the influence of various parameters on AR, CR, and WR, we define the following relation for the sensitivity:

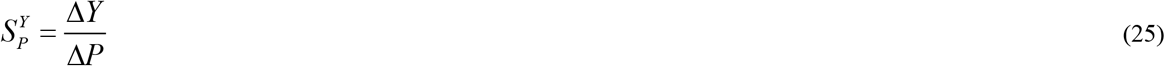

Here, it is supposed that initial values of M_1_ for all neurons of three cases in a 3×3 network is increased by 0.1 (i.e., ∆*M*_1_ = 0.1) and other parameters remain fixed unless otherwise stated. Fig. 8 shows the neuron states for new microglia values at *t* = 1 *year* for three cases. By comparing Fig. 8 (a) with Fig. 6 (a), it is concluded that for the first case 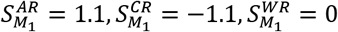. Therefore, the system is sensitive to microglia variations because the AR is increased from 0 to 11.1% (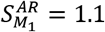, see neuron (3,3) in Fig. 8 (a), which has died). The numbers of red neurons have been decreased in this figure, compared to Fig. 6 (a). For the second case, the CR factor is increased (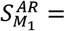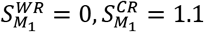). By comparing Fig. 6(c) with Fig. 8(c), one can see that the CR is increased while the WR is decreased for the third case 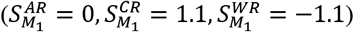.

**Fig. 8.**
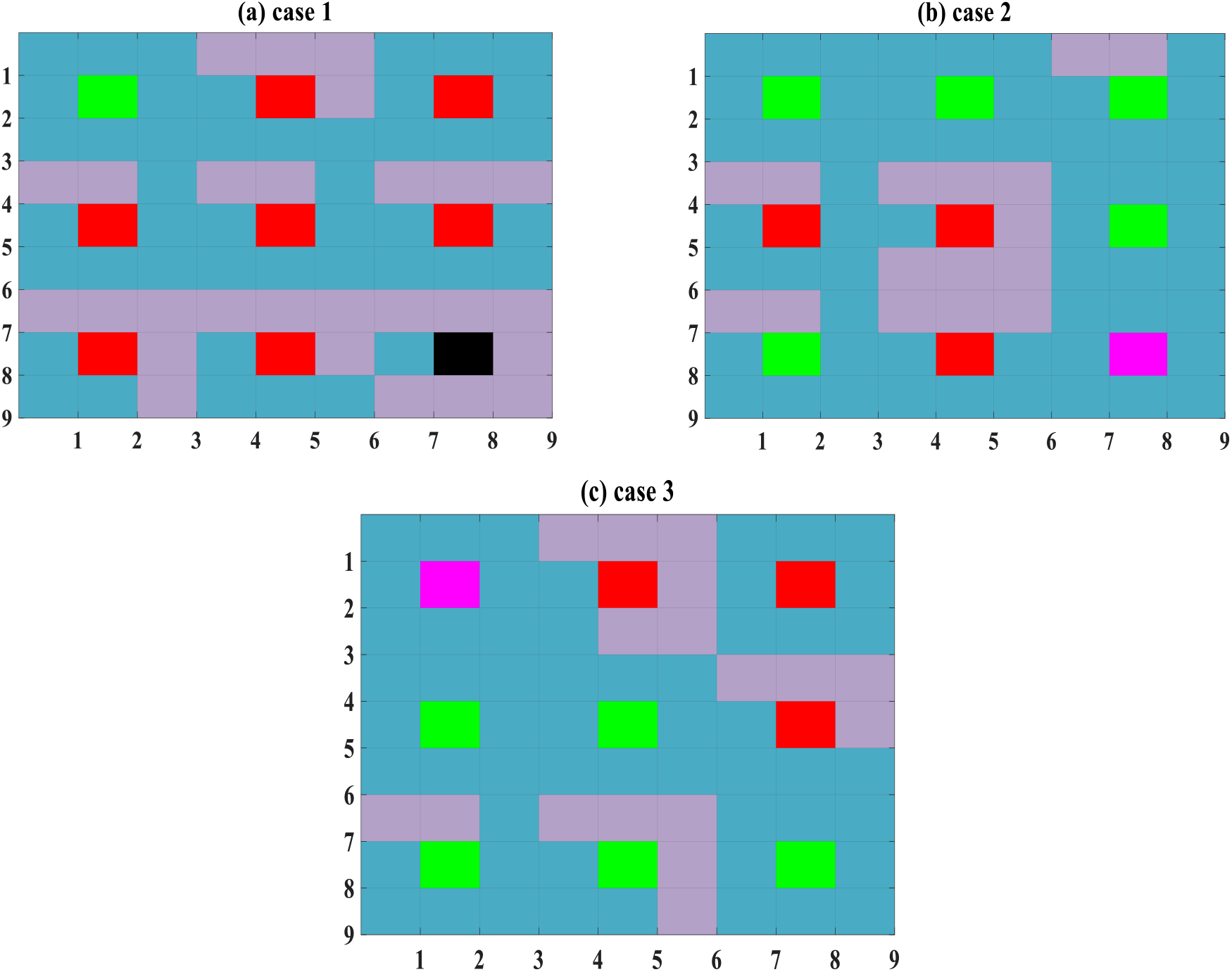
CA model of neurons and their synapses in a 3×3 network at *t* = 1 *year* for the sensitivity study of Microglia variations for three cases.

As a final point, the states of neurons have been represented in Fig. 9 to study the sensitivity of Astroglia variations. Again, we assume that initial values of A_P_ for all neurons of three cases in a 3×3 network is increased by 0.1 (i.e., ∆*A*_*p*_ = 0.1) and other parameters remain fixed unless otherwise stated. It is obvious that AR sensitivity for the first case 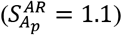 is similar to Fig. 8 (a) but the numbers of red neurons 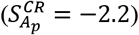 in Fig. 9 (a) is reduced more, compared to Fig. 8 (a). Hence, the sensitivity of neuron states to astroglia variations is much greater than microglia variations for the first case. For the second case, AR, CR, and WR are not sensitive to the Astroglia variations, i.e., 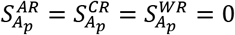 (compare Fig. 6(b) and 9(b)). It is obvious from Fig. 9(c) that the numbers of the red neurons are increased 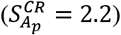 for the third case, compared to Fig. 8 (c). It can be concluded that the sensitivity of neuron states to astroglia variations is much greater than microglia variations for CR and WR factors, which confirms the results of (Thuraisingham 2017).

**Fig. 9.**
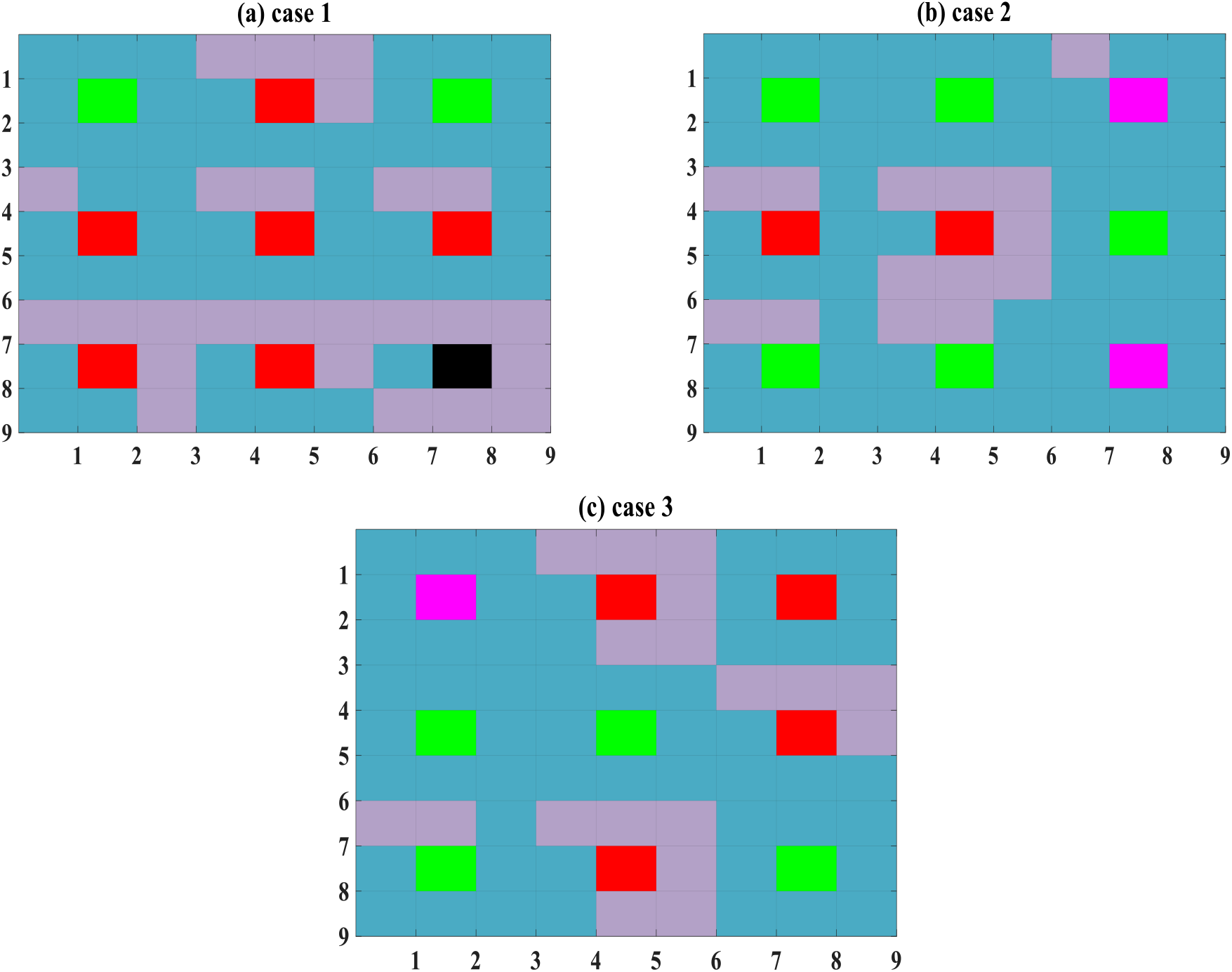
CA model of neurons and their synapses in a 3×3 network at *t* = 1 *year* for the sensitivity study of Astroglia variations for three cases.

## 4. Conclusion

In this paper, a novel mathematical model for the progress of Alzheimer’s disease was introduced and investigated. The basis of the model was the Cellular Automata network and differential equations of the Puri-Li article. The presented model studied schematically the neuron and its synapse states in a *M* × *L*-network. A new definition of AD rate was defined in this article. Moreover, two important factors, i.e., Critical and Warning Rates, were introduced here to determine the emergency state of AD progression. A 3×3 network was simulated in MATLAB software and studied in this paper. The proposed mathematical model could explain the internal mechanisms within the hippocampus very well. Our results showed that neuron states are much more sensitive to astroglia variations, in comparison to microglia variations. The presented study can assist the scientists to better and effectively predict, recognize and study the progression of AD.

## Ethical statement

This work is partially supported by Vice-Chancellor in Research Affair-Tabriz University of Medical Sciences under Ethical Code No. IR.TBZMED.VCR. REC.1399.377. The funders had no role in study design, analysis, numerical simulations, the decision to publish, or preparation of the manuscript.

